# Ring nucleases deactivate Type III CRISPR ribonucleases by degrading cyclic oligoadenylate

**DOI:** 10.1101/380436

**Authors:** Januka S Athukoralage, Christophe Rouillon, Shirley Graham, Sabine Grüschow, Malcolm F White

## Abstract

The CRISPR-Cas system provides adaptive immunity against mobile genetic elements in prokaryotes, utilising small CRISPR RNAs which direct effector complexes to degrade invading entities. Type III effector complexes were recently demonstrated to synthesise a novel second messenger, cyclic oligoadenylate (cOA), on binding target RNA. cOA in turn binds to and activates a range of downstream effector proteins including ribonucleases (Csm6/Csx1) and transcription factors via a CARF (CRISPR associated Rossman Fold) domain, inducing an antiviral state in the cell that is important for immunity. The mechanism of the “off-switch” that resets the system is not understood. Here, we report the identification of the nuclease that degrades these cOA ring molecules. The “Ring nuclease” is itself a CARF family protein with a metal independent mechanism, which cleaves cOA_4_ rings to generate linear di-adenylate species and switches off the antiviral state. The identification of Ring nucleases adds an important insight to the CRISPR-Cas system.

## Introduction

Cyclic oligoadenylate (cOA, with a ring size of 4 or 6 AMP monomers) has emerged as a key second messenger signalling the presence of invading mobile genetic elements (MGE) in prokaryotes harbouring type III (Csm/Cmr) CRISPR-Cas systems ^1, 2^. cOA is synthesised by the PALM/cyclase domain of the Cas10 subunit, which is activated by the binding of target RNA. cOA in turn activates a range of proteins with CARF (CRISPR associated Rossman Fold) domains including HEPN (Higher Eukaryotes and Prokaryotes Nucleotide binding) domain ribonucleases (Csm6/Csx1) and transcription factors ^3–5^, precipitating an antiviral state in infected cells that may enhance viral clearance, result in dormancy, or cell death ^6^. The Csm6 and Csx1 ribonucleases play an important role in CRISPR-based immunity in several systems ^7–9^. cOA is therefore a potent signalling molecule that must be tightly controlled if cells are to survive a viral infection. Synthesis of cOA is switched off when type III effectors cleave and release viral RNA targets ^1, 10^. However, this will not remove cOA already synthesised, potentially leading to untrammelled ribonuclease activity and cell death. A variety of other cyclic nucleotide signalling molecules, such as di-cAMP, which perform distinct signalling roles in prokaryotes, are degraded by specific phosphodiesterases ^11^, but the enzyme specific for cOA has not been identified. Here, we report the identification and characterisation of a family of enzymes that degrade cOA_4_, providing a mechanism for the deactivation of the antiviral state induced by cOA in cells harbouring a type III CRISPR-Cas system.

## Results

To identify the enzyme responsible for the degradation of cOA, we undertook a classical biochemical approach. We selected the crenarchaeal species *Sulfolobus solfataricus* for this study, as it has long been a model for biochemical studies of the type III CRISPR system. Starting with S. *solfataricus (Sso)* cell lysate, we noted the presence of an activity that converted radioactively labelled cOA_4_ into a form (X) that migrated more slowly in denaturing gel electrophoresis (Figure 1). We fractionated the cell lysate through three chromatography steps (phenyl-sepharose, size exclusion and heparin) and followed the activity. The final purification step was followed by assay of each fraction and SDS-PAGE analysis (Figure 1d), revealing that the activity correlated with a single band. This protein was identified by mass spectrometry as Sso2081 – a member of the CARF domain-containing protein family ^12^ (Figure S1). To confirm that Sso2081 was the nuclease responsible for degradation of cOA_4_, we expressed and purified the recombinant protein in *E. coli.* The enzyme converted cOA_4_ to a slower-migrating species on gel electrophoresis, as seen for the activity from S. *solfataricus* extracts, in a reaction independent of divalent metal ions, since activity is maximal in the presence of EDTA (Figure 1e). We noted that the Csx1 nuclease (Sso1389; PDB 2I71) ^10^ is found close to an uncharacterised CARF domain protein of known structure, Sso1393 (Figure S1) that is homologous to Sso2081 (Figure S2). Pure recombinant Sso1393 also exhibits cOA_4_ degradation activity, whilst Csx1 does not (Figure 1e). In contrast, of the three enzymes, only Csx1 displays cOA_4_-stimulated ribonuclease activity on a linear RNA substrate (Figure 1f). Sso2081 and Sso1393 degrade cOA_4_ with single-turnover rate constants of 0.23 ± 0.01 and 0.024 ± 0.0004 min^-1^, respectively (Figure 2a and S3). The ten-fold higher specific activity of Sso2081, combined with its higher expression levels in S. *solfataricus*^13^, suggest that it is the major cOA_4_-degrading enzyme in this organism.

**Figure 1.**
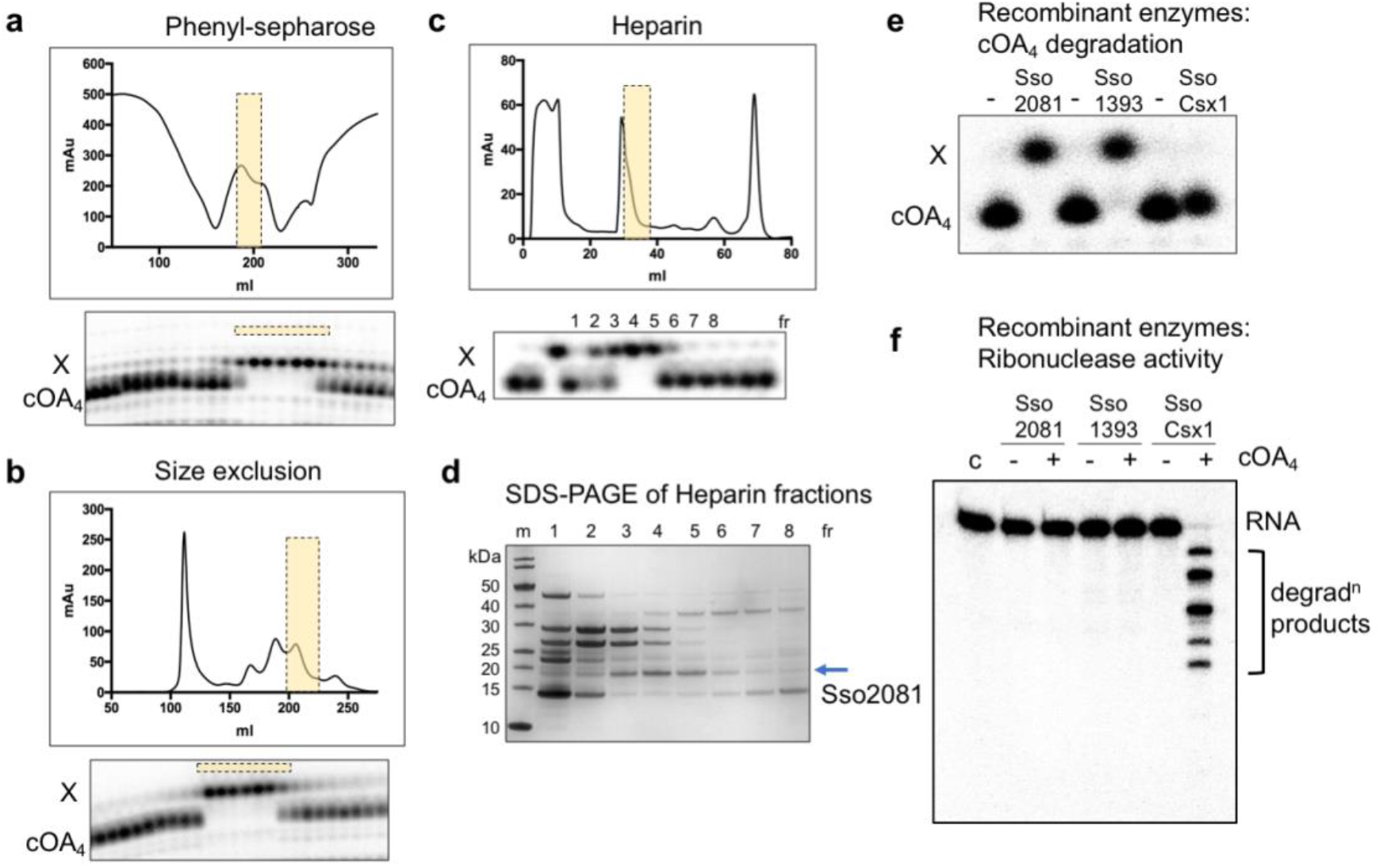
Purification and identification of the enzyme that degrades cOA_4_. A *S. solfataricus* cell lysate was fractionated by **a**, phenyl-sepharose, **b**, size exclusion and **c**, heparin chromatography. At each stage, fractions were assayed for cOA_4_ conversion activity using radioactive cOA_4_, and active fractions generating product X (indicated by shaded boxes) were pooled for the next stage. **d**, Following the heparin column, each fraction was assayed and analysed by SDS-PAGE. The band corresponding to the peak of activity (arrowed) was excised from the gel and identified by mass spectrometry as Sso2081, a CARF domain protein. **e**, Purified recombinant Sso2081 and 1393 degrade cOA_4_, but Csx1 does not. **f**, Only Csx1 degrades linear RNA in the presence of cOA_4_.

**Figure S1.**
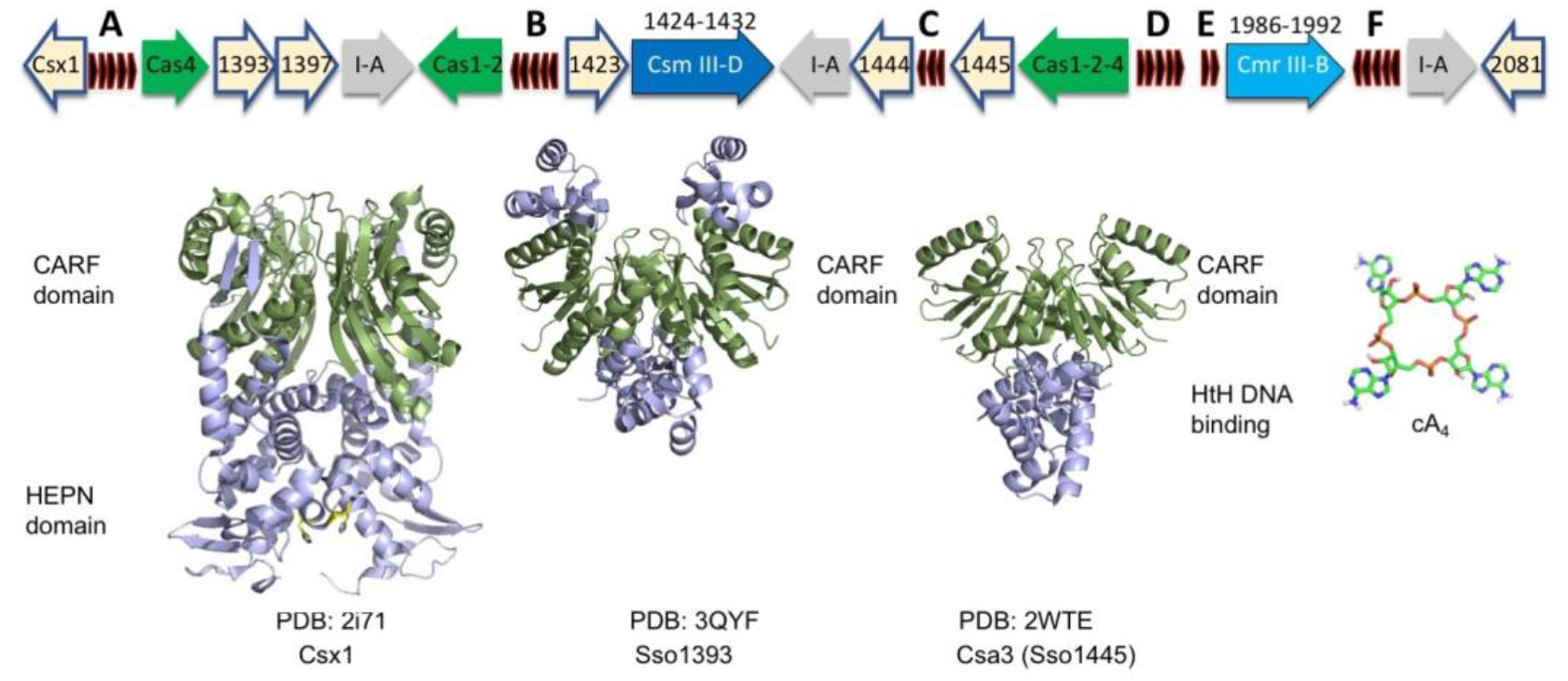
Genome organisation of the CRISPR-Cas locus of *S. solfataricus*. The type I-A, III-B and III-D effector complex operons are depicted, along with genes encoding adaptation proteins and the position of the six CRISPR loci (A-F). Genes outlined in blue encode proteins with CARF domains. The structures of three CARF family proteins are shown with CARF domains coloured green, along with the structure of cOA_4_.

Metal independent ribonucleases and ribozymes, including the Cas6 nuclease family ^14^, share a common mechanism: activation of the 2’-hydroxyl of the ribose sugar as the nucleophile that attacks the phosphodiester bond targeted for cleavage, leading to products with a cyclic 2’,3’-phosphate and a 5’-hydroxyl moiety ^15^. To determine the degradation products of cOA_4_ treated with Sso2081 and 1393, we analysed radioactively labelled species by thin layer chromatography (TLC) (Figure 2b). Sso2081 converted cOA_4_ into a faster-migrating form (X) that could be phosphorylated by treatment with polynucleotide kinase (PNK), showing that it was a linear product with a 5’-OH group (lanes 2 and 3). For Sso1393 an intermediate product (Y) with a 5’-OH group that converted over time into the final product (X) was also observed (lanes 4-7). Lanes 8 and 9 show standards generated by cleavage of RNA oligonucleotides by the MazF ribonuclease ^10^ to generate the species P-OA_2_>P (5’-phospho-ApAp with a cyclic 2’,3’ phosphate) and P-OA_4_>P, respectively. The final products of Sso2081 and 1393 run at the same position as the P-OA_2_>P standard once phosphorylated by PNK, whilst the intermediate observed for Sso1393 runs at the same position as the P-A_4_>P standard.

**Figure 2.**
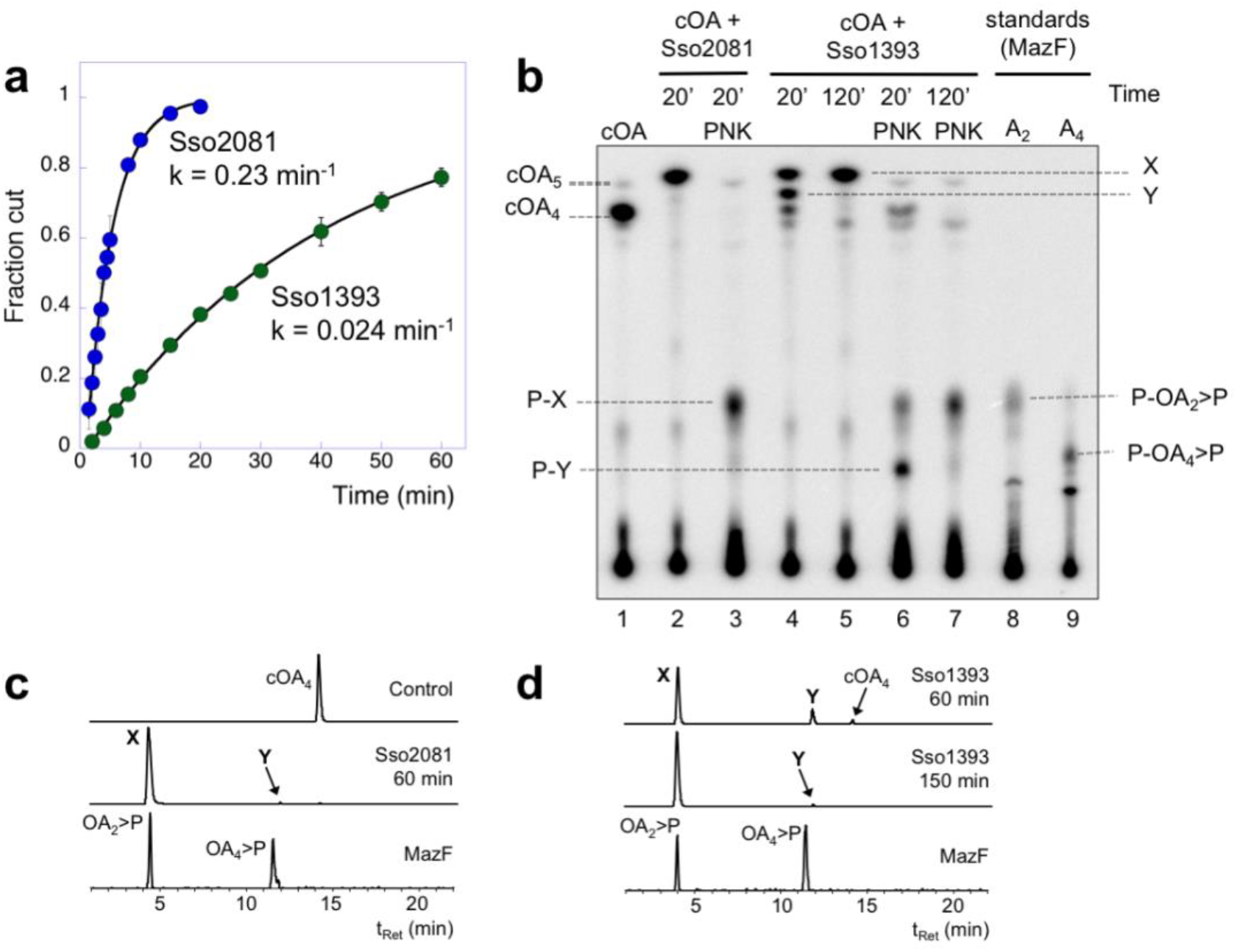
Kinetics and products of Ring nuclease activity. **a**, Single turnover kinetic analysis of cOA_4_ cleavage by Sso2081 and Sso1393. Sso2081 is a ten-fold faster enzyme. Data points are the means of triplicate measurements, with standard deviation shown. **b**, Thin layer chromatographic analysis of substrates and products. Lane 1 shows cOA_4_ synthesised by the Csm complex. Lane 2 shows the product (X) of conversion by Sso2081 after 20 min. Lane 3 shows product (X) following phosphorylation by PNK. Lanes 4-7 show the products (X and Y) of cOA_4_ conversion by Sso1393 after 20 and 120 min, respectively, before and after PNK treatment. Lanes 8 and 9 are markers for P-OA_2_>P and P-OA_4_>P generated by the MazF nuclease. **c, d**, LC-HRMS analysis of cOA_4_ cleavage by Sso2081 and Sso1393, respectively. Shown are ion chromatograms extracted for *m/z* 657.10 ± 0.5 corresponding to cOA_4_^-2^, OA_4_>P^-2^ and OA_2_>P^-1^. The control is cOA_4_ incubated for 150 min without enzyme. Traces for the conversion of cOA_4_ by Sso2081 after 60 min, and by Sso1393 after 60 and 150 min are shown. Products X and Y are indicated. The bottom traces show the linear oligoadenylates OA_2_>P and OA_4_>P generated by MazF as standards.

To identify products X and Y, we incubated Sso2081 and 1393 individually with cOA_4_ and analysed the reaction products by liquid chromatography-high resolution mass spectrometry (LC-HRMS, Figure 2c and d). Upon 60 min incubation with recombinant Sso2081, the peak corresponding to cOA_4_ had almost completely disappeared in favour of the main product (X) with a retention time of 4.4 min alongside a trace of compound Y eluting at 11.9 min (Figure 2c and S4). Linear OA_2_ and OA_4_ species with a 2’,3’-cyclic phosphate, generated using the MazF toxin ^10^, eluted with comparable retention times as products X and Y, respectively. Mass spectrometry revealed product X to have a neutral mass of 658.104 amu, consistent with linear A_2_>P (expected for C_20_H_24_N_10_O_12_P_2_ 658.105 amu, δm −1.6 ppm). The mass of product Y (1316.208 amu), on the other hand, was consistent with A_4_>P (expected for C_40_H_48_N_20_O_24_P_4_ 1316.210 amu, δm −1.6 ppm). The same products were observed for Sso1393 (Figure 2d and S4). Again, compound X was the main product with the peak corresponding to compound Y more visible at early time points compared to the Sso2081 reaction. The occurrence of one major product X for both enzymatic reactions and the detection of an apparent intermediate Y for the Sso1393 reaction are both in agreement with observations from TLC analysis, which demonstrates that product Y is only ever observed in very small amounts in the Sso2081 reaction, probably due to the significantly faster catalytic rate of this enzyme (Figure S5). Thus, these enzymes break the cOA_4_ ring using a metal-independent mechanism, generating linear A_4_>P intermediate and A_2_>P products with 5’-OH and 2’,3’-cyclic phosphate termini. We propose the collective term “Ring nucleases” for this family of phosphodiesterases.

**Figure S2.**
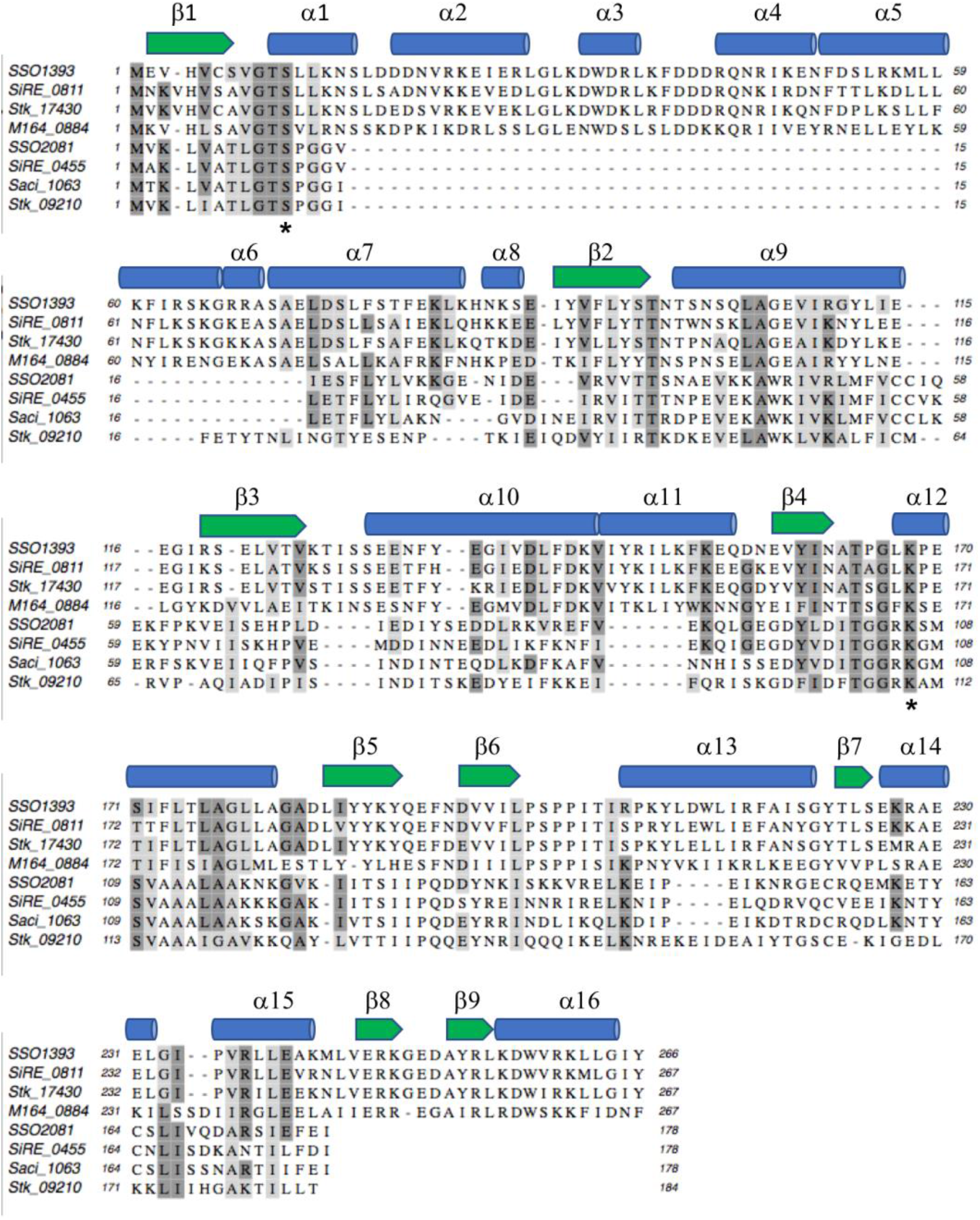
Structure-guided sequence alignment of Sso1393, Sso2081 and orthologues. Multiple sequence alignment showing Sso1393 with three homologues from the *Sulfolobales* aligned with Sso2081 and 3 homologues. Secondary structure is shown above the alignment, based on the structure of Sso1393 (PDB 3QYF). Ser-11 and Lys-168 are indicated by asterisks. Conserved residues are shaded. Sequences aligned are from *S. solfataricus* (Sso1393, Sso2081); *S. islandicus* REY15A (SiRE_0811 SiRE_0455); S. islandicus M.16.4 (M164_0884); *S. acidocaldarius* (Saci_1063) and *S. tokodaii* (Stk_17430).

The structure of Sso1393 (PDB 3QYF, unpublished) reveals a canonical CARF domain formed by a homodimeric subunit arrangement, with a C-terminal extension (Figure S1). To help understand the mechanism of Ring nucleases, we docked cOA_4_ into the CARF domain of Sso1393 (Figure 3a and S6). This is a simple model without any attempt at energy minimisation but allows hypotheses concerning the enzyme mechanism to be drawn. Using structure-guided multiple sequence alignment of Sso1393, Sso2081 and homologues, we identified conserved residues positioned close to the cOA_4_ binding site that might play a role in catalysis (Figure 3, S2, S6). We noted the conservation of a lysine (K168 in Sso1393) that is predicted to lie in the centre of the cOA_4_ binding site and which could thus play a role in catalysis. Sso2081 has two basic residues (R105/K106) in this position. The K168A variant of Sso1393 was catalytically inactive, as was the Sso2081 R105A/K106A variant, confirming an important role in cOA catalysis, possibly by stabilising the pentacovalent phosphorous generated in the transition state ^15^ (Figure 3b, S6d). The absolutely conserved residue S11 (Figure 3), was also targeted by mutagenesis. The S11A variant had a significantly lower catalytic rate constant for both enzymes, with kcat reduced 3.5 and 32-fold for Sso2081 and Sso1393, respectively (Figure 3b, S6). Consistent with the absence of Ring nuclease activity, Csx1 lacks these key active site residues.

**Figure 3.**
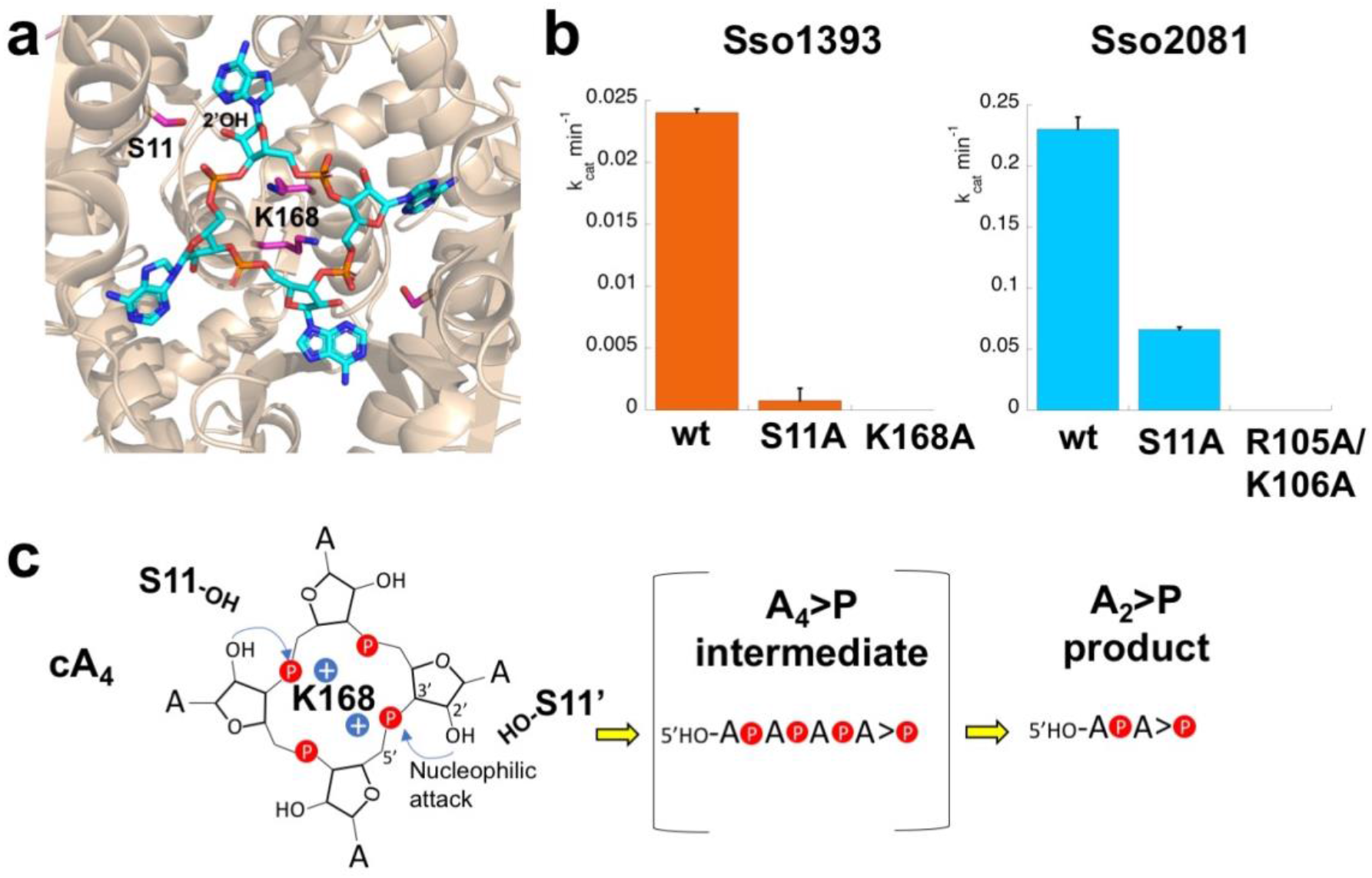
Structure and mechanism of Ring nucleases. **a**, Structure of the CARF domain of Sso1393 with cOA_4_ docked. The active site residues K168 and S11 are shown. **b**, Kinetic analysis of Sso1393 (wild-type, S11A and K168A variants) and Sso2081 (wild-type, S11A and R105A/K106A variants). Catalytic rate constants under single turnover conditions are plotted. The data are derived from triplicate rate measurements (Figure S6d), and standard errors are shown. **c**, Cartoon showing the reaction scheme for conversion of cyclic to linear OA_4_ and OA_2_. S11 may participate in the correct positioning of the 2’-OH group of the ribose to facilitate nucleophilic attack, whilst the basic residue K168 (and R105/K106 in Sso2081) is essential for catalysis and may stabilise the pentacovalent phosphorus formed in the transition state.

Based on the evidence available, we propose that cleavage of cOA_4_ is catalysed by binding the molecule in an orientation that allows the 2’-hydroxyl of the ribose to attack the bridging phosphorus, coupled with stabilisation of the developing transition state by the basic residue(s) at the base of the binding pocket (Figure 3c). The single turnover rate constant for Sso2081, at below 1 min^-1^, is comparable to the Cas6 nuclease family, which use a similar mechanism ^14, 16^. Given the dimeric organisation of the CARF domain there is the possibility for two active sites acting on opposite sides of the cOA_4_ ring, consistent with the observation of an OA_2_>P product. For the slower Sso1393 enzyme, appreciable levels of OA_4_>P intermediate were observed, suggesting that the two active sites need not function in a concerted manner.

We next wished to reconstitute the cOA signalling system *in vitro* to confirm the function of the Ring nucleases. We set up a reaction containing the Csm (Type III-D) effector complex, 0.5 mM ATP and a variable concentration of target RNA from 10 to 0.01 nM, to activate cOA_4_ synthesis. After incubation for 1 h at 70 °C in the presence or absence of Sso2081, the HEPN ribonuclease Csx1 together with radioactively labelled Csx1 substrate RNA were added. Target RNA binding by Csm activates the Cyclase domain, switching on cOA production. The Cyclase domain is deactivated when target RNA is cleaved and dissociates from Csm, leaving the complex free to bind to further targets. Higher target RNA concentrations thus result in higher levels of cOA_4_ ^10^ and stronger activation of Csx1, which degrades a labelled substrate RNA (Figure 4a). The presence of Sso2081 abrogated the activation of Csx1 in a target RNA concentration-dependent manner. This recapitulates the situation likely to prevail in cells infected with a virus, where viral RNA load determines the extent of cOA production and speed of subsequent clearance by the Ring nuclease (Figure 4b).

**Figure 4.**
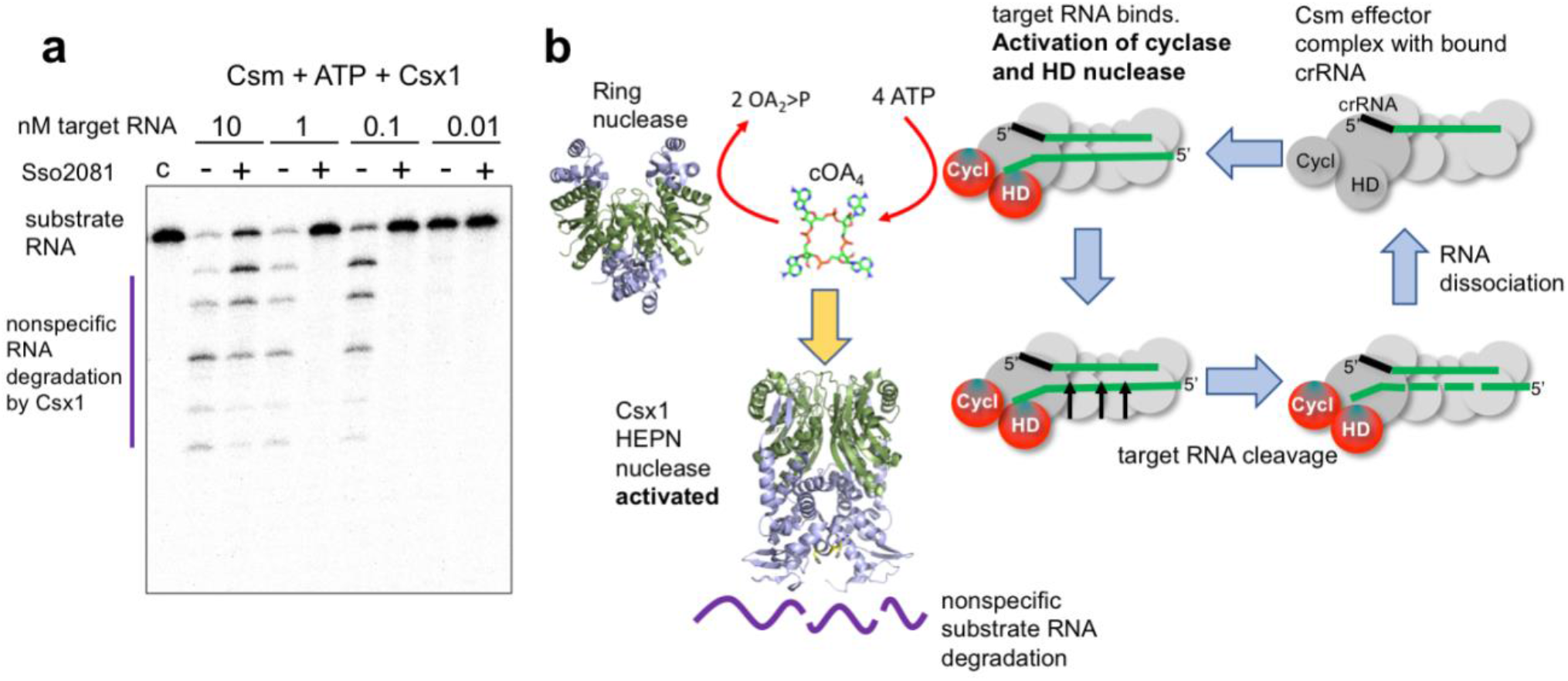
Reconstitution of the cOA signalling pathway. **a**, The Csm effector generates cOA_4_ in proportion to the amount of viral target RNA present ^10^, activating the HEPN nuclease Csx1. The presence of Sso2081 (2.5 μM) for 1 h prior to the addition of Csx1 (0.5 μM) for 20 min at 70 °C reversed Csx1 activation partially when 10 nM target RNA was present, and fully when lower amounts of RNA were used. Control c shows the experiment in the presence of Sso2081 and absence of target RNA. **b**, Schematic of the cOA signalling system. Type III (Csm/Cmr) effector complexes loaded with crRNA bind to cognate viral target RNA, activating the cyclase and HD nuclease domains. cOA is synthesised, which in turn activates HEPN nucleases such as Csx1. Target RNA cleavage and subsequent dissociation switches off the Cyclase domain, stopping synthesis of cOA. The Ring nucleases complete the deactivation of the antiviral state by degrading extant cOA.

Thus, we have identified a family of CARF-domain containing Ring nucleases as the enzymes responsible for the degradation of cyclic oligoadenylate, which is synthesised by type III CRISPR systems in response to detection of viral RNA. The Ring nucleases can be considered to act as the “off-switch” for the system, limiting the damage caused by the HEPN ribonucleases activated by cOA once invading RNA has been cleared from the cell. Many type III CRISPR-Cas systems have multiple associated CARF-domain containing proteins, some of which may be specialised for the degradation of cOA. In a minimal system, it is possible that a single enzyme has a C-terminal HEPN family ribonuclease coupled to an N-terminal CARF family Ring nuclease. This would allow cOA binding to rapidly switch on the ribonuclease activity and then slowly auto-deactivate by cOA cleavage. However, specialisation of CARF proteins as Ring nucleases would yield the advantage of allowing levels of cOA-activated ribonuclease and cOA-degrading Ring nuclease activity to be controlled independently. It is also possible that cOA_6_ rings such as those generated in *Streptococcus thermophilus* will be processed differently, as they are likely to have more conformational flexibility than cOA_4_. The recent identification of a wide range of as-yet uncharacterised CARF domain proteins across the prokarya ^5^ suggests that we have only scratched the surface of this system. These are fruitful areas for future studies.

## Acknowledgements

This work was funded by grants from the Biotechnology and Biological Sciences Research Council (REF BB/M000400/1 and BB/M021017/1). MFW is a Wolfson Research Merit Award holder. We acknowledge the contribution of the Mass Spectrometry Unit of the University of St Andrews to this work.

## Author contributions

J.S.A and C.R. carried out and analysed the bulk of the experimental work; S.Grü. carried out and analysed the mass spectrometry; S.Gra. generated expression plasmids and purified proteins; M.F.W. oversaw the work, analysed the data and wrote the manuscript. All authors contributed to the data analysis and writing.

---

**Figure S3.**
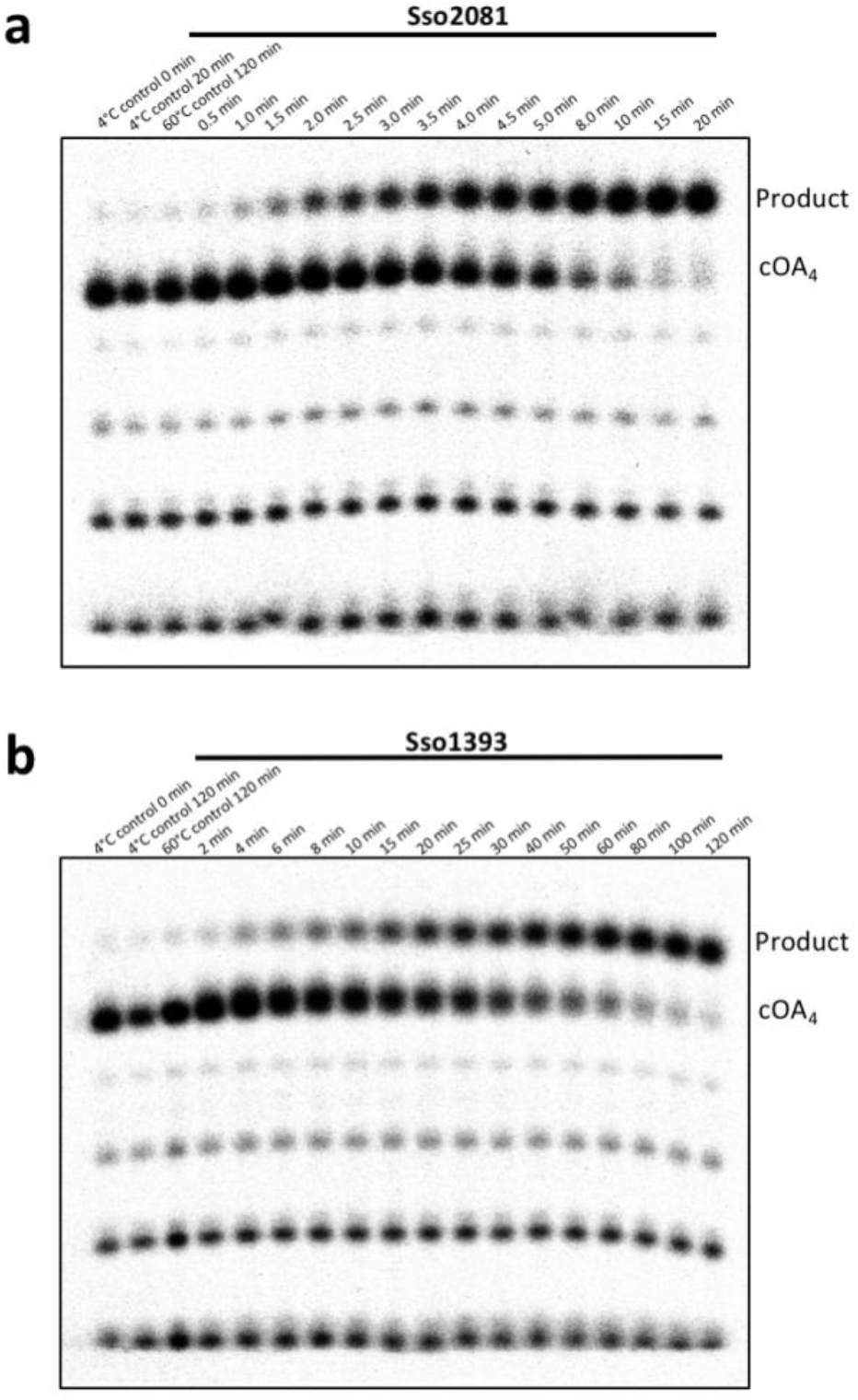
Single-turnover kinetic analysis of cOA_4_ degradation by a, Sso2081 and b, Sso1393. Representative phosphorimages of denaturing polyacrylamide gel electrophoresis (PAGE) analysing the reaction of radioactively labelled cOA_4_ with **a**, 2 μM Sso2081 or **b**, Sso1393 at 60 °C over time. Triplicate experiments were carried out, quantified by phosphorimaging and the fraction of cOA_4_ cleaved plotted in Figure 3b.

**Figure S4.**
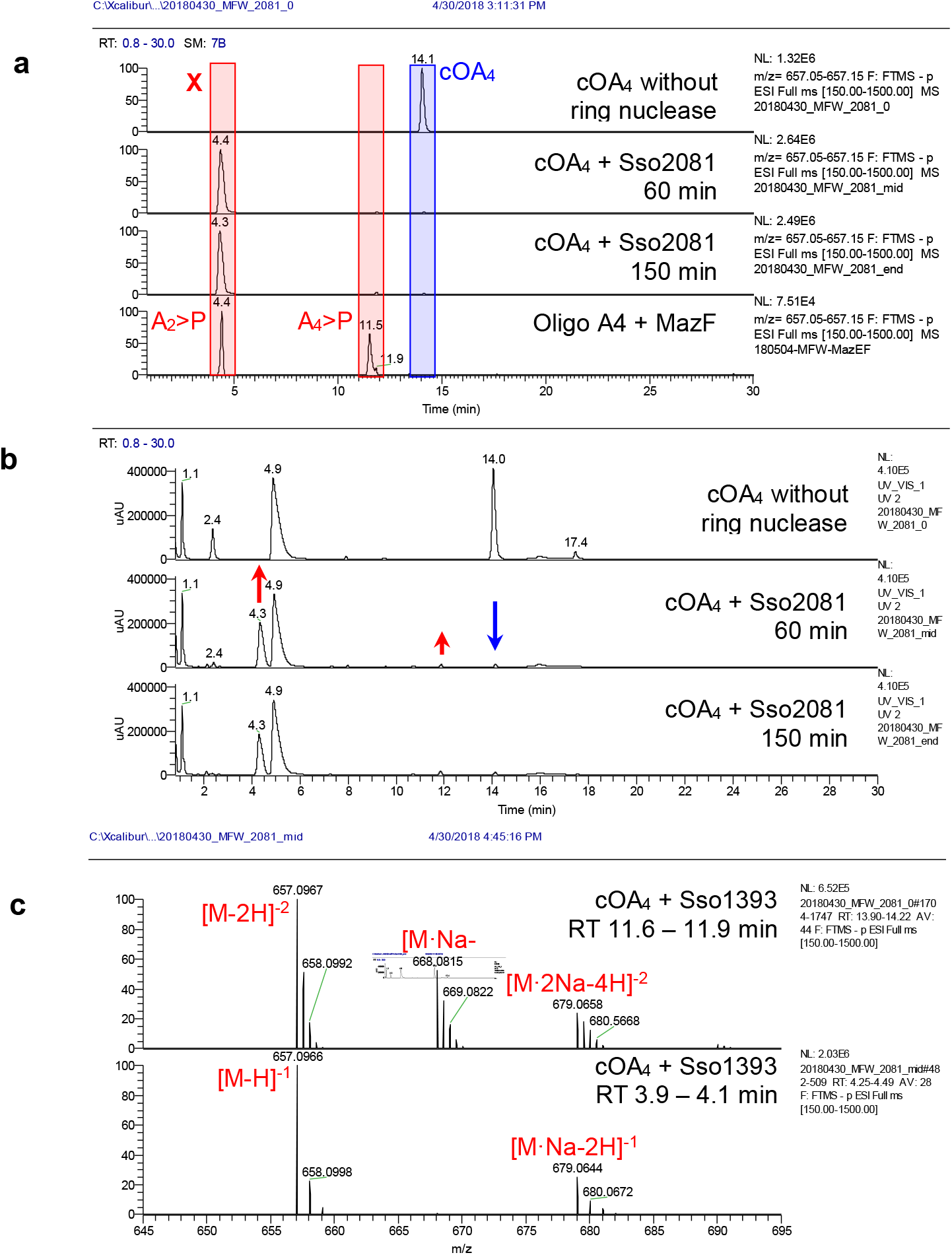
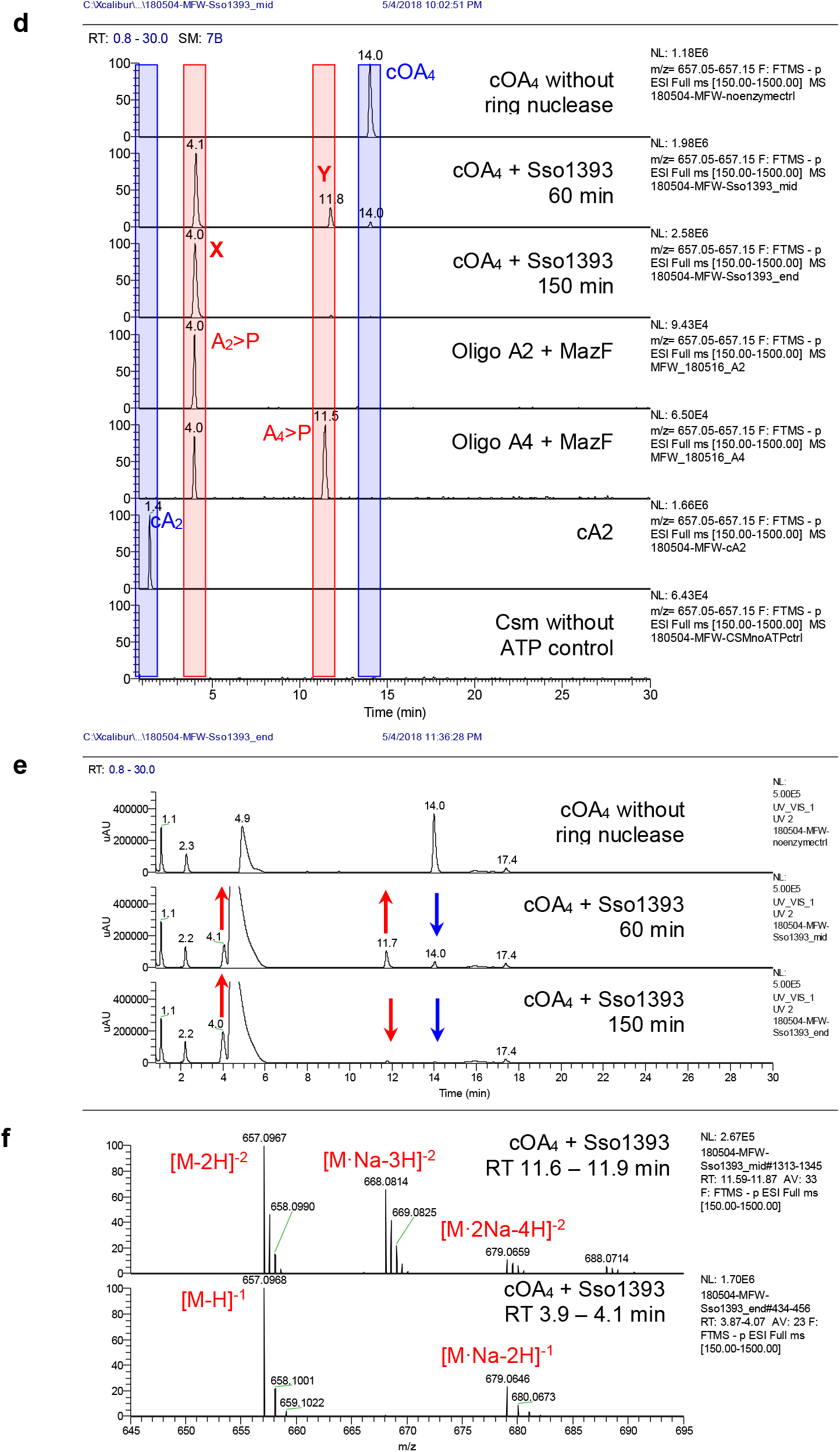
LC-MS analysis of Sso1393 (A-C) and Sso2081 (D-F) reactions. **a**, Ion chromatograms extracted for *m/z* 657.1 (cA_2_^-1^, A_2_>P^-1^, cA_4_^-2^, A_4_>P^-2^). Individual lanes are labeled. cOAs (mainly cOA_4_), derived from reaction of Csm with ATP, was incubated with Sso1393 for 60 and 150 min or without ring nuclease for 150 min. A control in which the ATP from the Csm cyclase reaction had been omitted was also analysed as control. Linear oligoadenylates with 2’,3’-cyclic phosphate were derived from hydrolysis of suitable DNA oligonucleotide subtrates with the toxin MazF; whereas cA_2_ was a commercially available standard. The traces show clearly the difference in retention time between the linear and cyclic isomers. **b**, UV traces at 254 nm. Peaks that change in intensity over the course of the enzymatic reaction are indicated by arrows. The three peaks that decreased or increased over the course of the reaction all match the changes in abundance of the *m/z* 657.1 species. The broad peak at 4.1 – 6 min is an unknown contaminant likely resulting from the phenol-chloroform extraction. **c**, Mass spectra for species X and Y. Calculated for C_20_H_23_N_10_O_12_P_2_^-1^ (cA_2_ / A_2_>P) *m/z* 657.0978, found 657.0968 (ám −1.4 ppm); calculated for C_40_H_46_N_20_O_24_P4^-2^ (cA_4_ / A_4_>P) *m/z* 657.0978, found 657.0967 (ám – 1.6 ppm). **d**, Ion chromatograms extracted for *m/z* 657.1 (cA_2_^-1^, A_2_>P^-1^, cA_4_^-2^, A_4_>P^-2^). Individual lanes are labeled. cOAs (mainly cOA_4_), derived from reaction of Csm with ATP, was incubated with Sso2081 for 60 and 150 min. Linear oligoadenylates with 2’,3’-cyclic phosphate were derived from hydrolysis of A4 RNA oligonucleotide with MazF. **e**, UV traces at 254 nm. Peaks that change in intensity over the course of the enzymatic reaction are indicated by arrows. The three peaks that decreased or increased over the course of the reaction all match the changes in abundance of the *m/z* 657.1 species. No changes are observed after 60 min reaction time. The broad peak at 4.9 min is an unknown contaminant likely resulting from the phenol-chloroform extraction. Shifts in retention time are possibly due to matrix effects. **f**, Mass spectra for cOA_4_ and product X. Calculated for C_20_H_23_N_10_O_12_P_2_^-1^ (cA_2_ / A_2_>P) *m/z* 657.0978, found 657.0966 (ám −1.8 ppm); calculated for C_40_H_46_N_20_O_24_P_4_^-2^ (cA_4_ / A_4_>P) *m/z* 657.0978, found 657.0967 (ám – 1.6 ppm).

**Figure S5.**
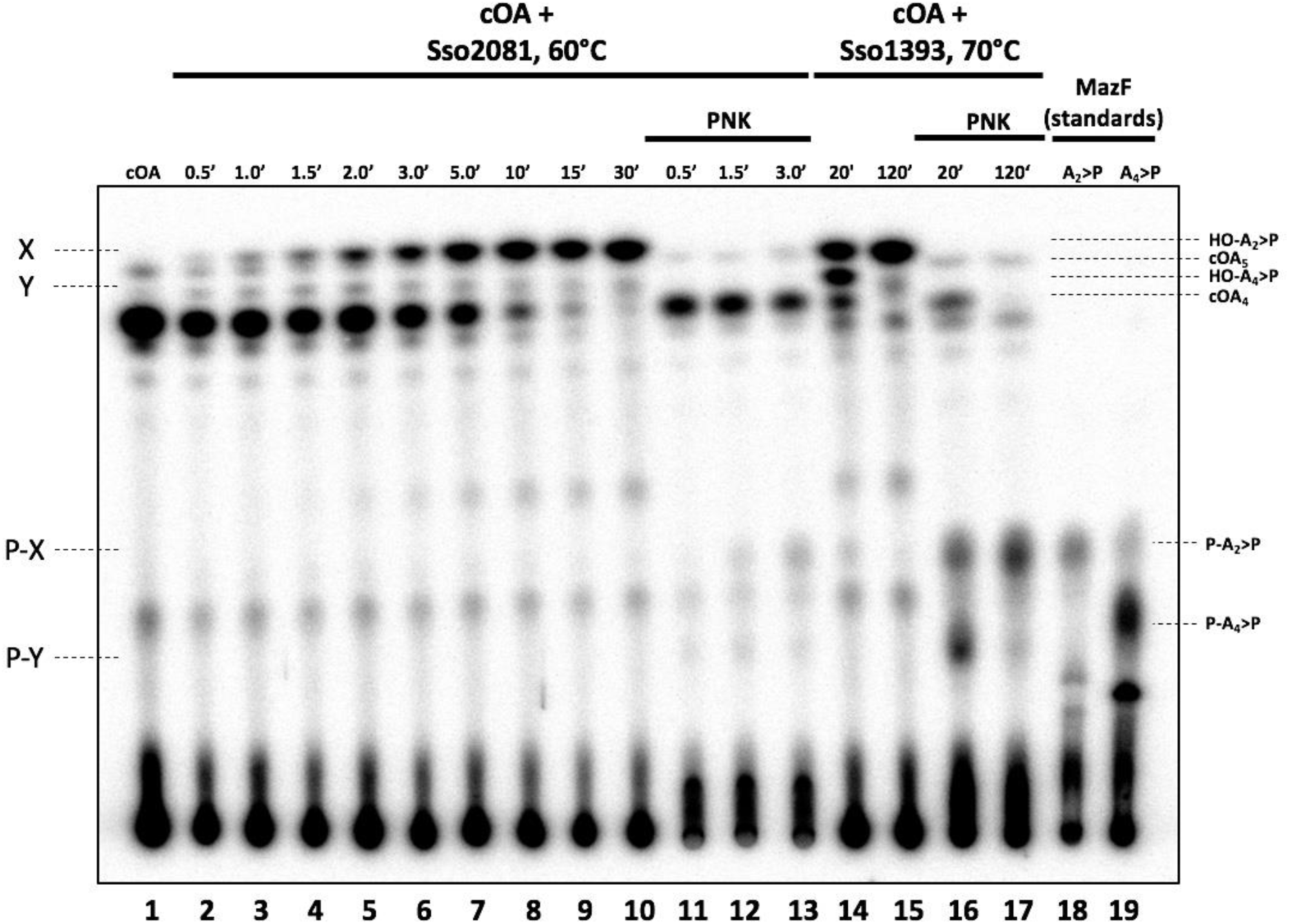
Sso2081 and Sso1393 cOA_4_ degradation mechanism investigated by TLC. Csm generated cOA (lane 1) was incubated with 2 μM Sso2081 dimer at 60 °C to determine the intermediate (Y) and final (X) reaction product over time (lanes 2-10). Lanes 11-13 show reaction product seen in lanes 2, 4 and 6 5’-end phosphorylated using T4 polynucleotide kinase (PNK) for identification of reaction intermediates and products by comparison to 5’-end phosphorylated mazF nuclease generated HO-A_2_>P and HO-A_4_>P standards. Lanes 14 & 15 show the reaction products of 2 μM Sso1393 dimer incubated with cOA at 70 °C for 20 and 120 min, respectively. Reaction product from lanes 14 & 15 are 5’-end phosphorylated by PNK for comparison to P-A_2_>P and P-A_4_>P standards. Comparison of PNK treated reaction product to standards showed the presence of a low amount of intermediate (P-Y) during the Sso2081 cOA_4_ cleavage reaction, which migrated similarly to the P-A_4_>P standard and did not change in abundance over time, whereas the abundance of the final product (P-X) increased over time. In contrast, comparison of Sso1393 PNK treated 20 min and 120 min reaction products showed a decrease of the intermediate (P-Y) over time and increase of product (P-X). A small amount of cOA5 was also present in Csm cOA, migrating between products X and Y, and consistent with its cyclic nature remained unchanged by PNK treatment.

**Figure S6.**
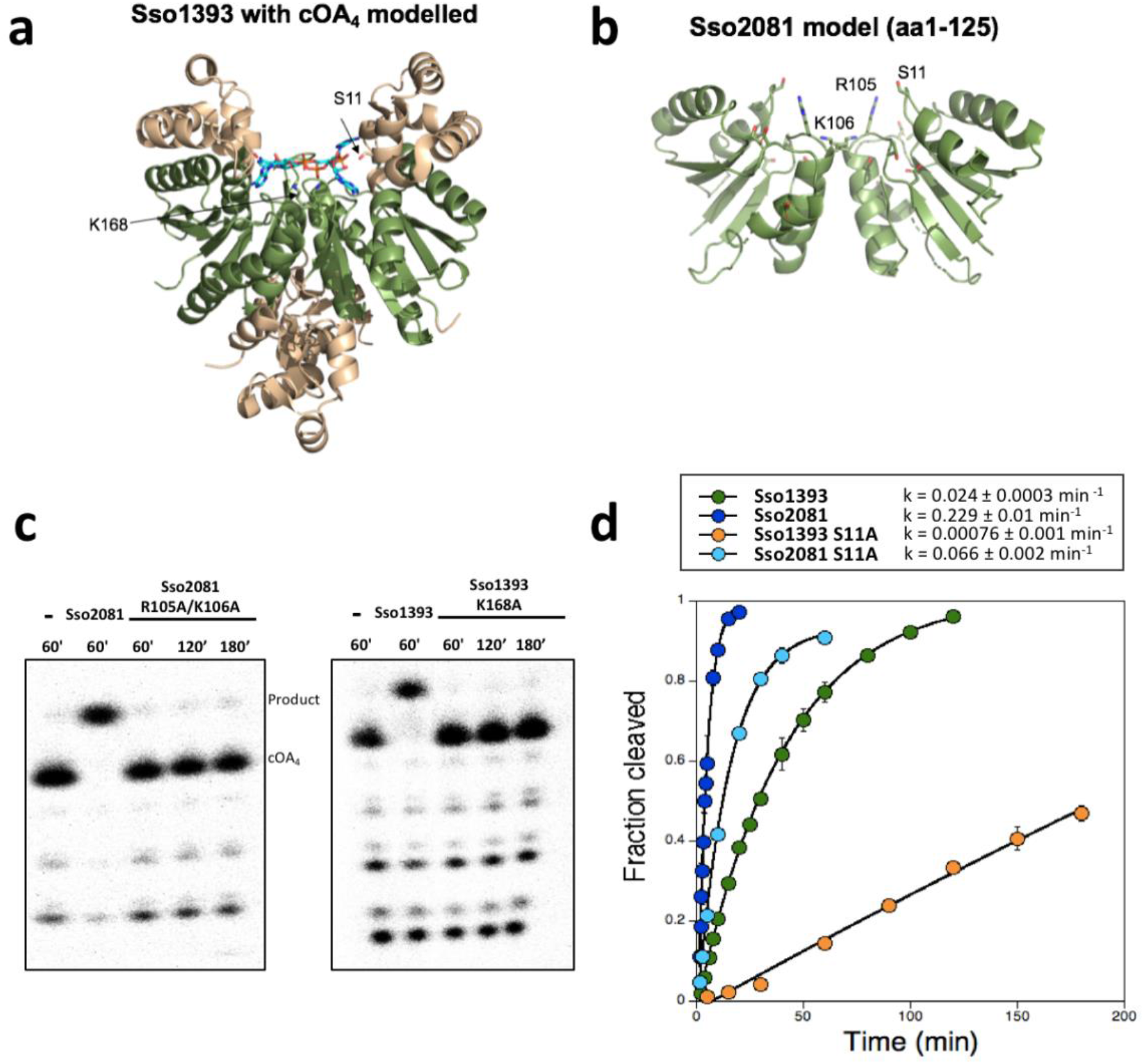
Structure and kinetic analysis of Sso1393 and Sso2081. **a**, Structure of Sso1393 (PDB 3QYF), with cOA_4_ docked at the active site. This is an orthogonal view to that shown in Figure 3. The CARF domain is coloured green. The side chains of Ser-11 and Lys-168 are shown. **b**, Model of the Sso2081 structure. Only the CARF domain (aa1-125) is modelled. Conserved residues Ser-11, Arg-105 and Lys-106 are labelled. Model generated using Phyre ^1^. **c**, Representative phosphorimages of denaturing PAGE assessing the cOA_4_ degradation activity of Sso2081 compared to its catalytically inactive R105A/K106A mutant over time (left hand panel), and Sso1393 activity compared to its K168A mutant (right hand panel). All reactions were carried out at 70 °C with 2 μM protein dimer. **d**, Single-turnover kinetics of Sso2081 and Sso1393 plotted alongside their S11A active site mutants and fitted to an exponential equation. Rate constants (fraction of cOA_4_ cleaved per min) are displayed with the legend (Sso2081; 0.23 ± 0.01 min^-1^, Sso2081 S11A; 0.066 ± 0.002 min^-1^, Sso1393; 0.024 ± 0.0003 min^-1^, Sso1393 S11A; 0.00076 ± 0.001 min^-1^). Experiments were carried out in triplicate and error bars show standard deviation.

## Materials and Methods

### Purification of cOA degrading enzyme from S. solfataricus cellular extract

*S. solfataricus (Sso)* P2 was grown as previously described ^17^ and cells pelleted by centrifugation at 4000 rpm (Beckman Coulter Avanti JXN-26; JLA8.1 rotor) for 15 min at 4 °C. Cells were suspended in buffer A containing 100 mM sodium phosphate pH 7.0 and 1.5 M ammonium sulphate with one EDTA-free protease inhibitor tablet (Roche). Cells were lysed by sonicating six times for 1 min with 1 min rest intervals on ice, and the lysate was ultracentrifuged at 40000 rpm (Beckman Coulter Optima L-90K; 70 Ti rotor) and 4 °C for 45 min before filtering and loading onto a phenyl-sepharose column (GE Healthcare) pre-equilibrated with buffer A. Protein was eluted with a linear gradient of buffer B containing 100 mM sodium phosphate pH 7.0 across 10 column volumes (CV). Each fraction was assayed for cyclic tetra-adenosine (cOA_4_) degradation activity and fractions displaying cOA_4_ degradation were pooled and concentrated using a 10000 MWCO ultracentrifugal concentrator (Amicon Millipore). Concentrated protein was then further separated by size-exclusion chromatography (S200; 26/60 GE Healthcare) in buffer containing 20 mM 4-(2-hydroxyethyl)-1-piperazineethanesulfonic acid) (HEPES) pH 7.5 and 150 mM KCl. Fractions were assayed for cOA_4_ degradation, and active fractions pooled and concentrated as previously, exchanging into 10 mM 2-(N-morpholino)ethanesulfonic acid (MES) pH 6.0 buffer during concentration. Concentrated protein was then loaded on to a 5 ml HiTrap heparin column (GE Healthcare) equilibrated with 10 mM MES pH 6.0 buffer, washing unbound protein with 8 % buffer C containing 10 mM MES pH 6.0 and 1.0 M NaCl for 2 CV. Protein was eluted by a linear gradient to 30 % buffer C across 5 CV followed by a gradient to 100 % buffer C across 4 CV. After assaying for cOA_4_ degradation, fractions of interest were visualised by sodium-dodecyl-sulfate polyacrylamide gel electrophoresis (SDS-PAGE). A protein band of ~18 kDa, the presence and abundance of which corresponded to a peak in cOA_4_ degradation was excised, trypsin digested and identified by mass spectrometry.

### Cloning, expression and purification of Sso2081, Sso1393 and variants

The cloning, expression and purification of Sso1389 (Csx1) has been described previously ^10^. A synthetic gene encoding Sso2081 was purchased from Integrated DNA Technologies, Coralville, IA, United States (IDT), while Sso1393 was PCR amplified from *S. solfataricus* P2 genomic DNA. Genes encoding Sso2081 and Sso1393 were cloned into the pEHisTEV vector ^18^, and transformed into *E. coli* DH5α competent cells. Sso2081 S11A, Sso2081 R105A/K106A, Sso1393 S11A and Sso1393 K168A variants were generated by site directed mutagenesis using the QuickChange Site-Directed Mutagenesis protocol (Agilent Technologies), with DNA primers purchased from IDT. Sequence verified constructs were transformed into C43 (DE3) *E. coli* cells for protein expression.

Expression of recombinant Sso2081 and variants was induced with 0.4 mM isopropyl β-D-1-thiogalactopyranoside at an OD_600_ of ~0.8, and cells incubated at 16 °C overnight with shaking at 180 rpm before harvesting by centrifugation at 4000 rpm (JLA8.1 rotor) at 4 °C for 15 minutes. Cells were suspended in lysis buffer (50 mM Tris-HCl pH 8.0, 0.5 M NaCl, 10 mM imidazole and 10% glycerol) with 1 mg/ml chicken egg lysozyme (Sigma-Aldrich) and one EDTA-free protease inhibitor tablet. Cells were sonicated six times for 1 min with 1 min rest intervals on ice at 4 °C. Cell lysate was then ultracentrifuged at 40000 rpm (70 Ti rotor) for 45 min at 4 °C, filtered and loaded onto a 5 ml HisTrap FF column (GE Healthcare) equilibrated with buffer E containing 50 mM Tris-HCl pH 8.0, 0.5 M NaCl, 30 mM imidazole and 10% glycerol. After washing unbound protein with 20 CV buffer E, recombinant Sso2081 and mutants were eluted with a linear gradient of buffer E supplemented with 0.5 M imidazole across 15 CV, then holding at 20% buffer F for 4 CV. Pooled fractions were concentrated as described previously, and the hexa-histidine affinity tag was removed by incubating protein with Tobacco etch virus (TEV) protease (10:1) for 4 h at 37 °C. His-tag cleaved Sso2081 was isolated from TEV by affinity chromatography as detailed above, eluting with buffer E prior to further purification by size-exclusion chromatography (S200 26/60; GE Healthcare) in buffer F (20 mM Tris-HCl pH 8.0, 0.5 M NaCl and 1 mM DTT).

Expression of recombinant Sso1393 (wild-type and variants) was induced as above, except cells were grown at 16 °C overnight before harvest. Cells were lysed and protein purified as for Sso2081. All proteins were aliquoted, flash frozen with liquid nitrogen, and stored at −80 °C.

### Cyclic oligoadenosine nuclease assays and kinetic analysis

^32^P-α-ATP incorporated cyclic oligoadenylate was generated using the Csm complex as previously described ^10^. Briefly, the Csm complex was incubated for 2 h at 70 °C in 100 μl final reaction volume in a pH 5.5 buffer containing 20 mM MES, 100 mM potassium glutamate and 1 mM DTT supplemented with 2 mM MgCl_2_, 1 mM ATP, 3 nM^32^P-α-ATP and 100 nM A_2_6 target RNA (5’-AGGGUCGUUGUUAAGAACGACGUUGUUAGAAGUUGGGUAUGGUGGAGA). The reaction was stopped by phenol-chloroform extraction followed by chloroform extraction. Sso1393, Sso2081, and their mutants were assayed for labelled cOA_4_ degradation by incubating 2 μM protein dimer with 1/300 diluted Csm generated^32^P labelled cOA (0.33 μl per 100 μl reaction) in buffer G (20 mM 2-amino-2-(hydroxymethyl)-1,3-propanediol pH 8.0, 100 mM NaCl, 1 mM EDTA and 1 mM DTT) at 70 °C. At desired time points, a 10 μl aliquot was removed, and the reaction quenched by adding to chilled phenol-chloroform (Ambion). Subsequently, 5 μl of deproteinised reaction product was extracted into 5 μl 100% formamide for denaturing PAGE. Control reactions include cOA incubated in buffer G without protein at 4 °C and at 60 °C up to the endpoint of each experiment (typically 20, 120 or 180 min) phenol-chloroform extracted as above. All experiments were carried out in triplicate. cOA_4_ degradation was visualised by phosphorimaging following denaturing PAGE (7M Urea, 20% acrylamide, 1x TBE). For kinetic analysis, cOA_4_ cleavage was quantified using the Bio-Formats plugin ^19^ of ImageJ ^20^ as distributed in the Fiji package ^21^ and fitted to a single exponential curve using Kaleidagraph (Synergy Software), as described previously ^22^.

### HEPN nuclease deactivation assays

For degradation of cOA_4_, Csm generated cOA was incubated with 2 μM Sso2081 dimer or 4 μM Sso1393 dimer in buffer K for 60 min and 120 min, respectively. Subsequently, the reaction was deproteinised by phenol-chloroform extraction and diluted two-fold in RNase free water. As a control reaction, cOA was mock treated in buffer with water in place of protein and deproteinised as before. 500 nM Csx1 was incubated with 50 nM A1 RNA and either no cOA activator, 1/100 diluted untreated Csm cOA, 1/100 mock treated cOA or 1/100 ring nuclease treated cOA in buffer containing 20 mM MES pH 5.5, 100 mM K-glutamate and 1 mM DTT for 60 min at 70 °C. Reactions were quenched by the addition of a reaction volume equivalent of 100% formamide, and RNA cleavage was assessed by phosphorimaging following denaturing PAGE.

### Reconstitution of the cOA signalling pathway

The Csm complex (70 nM carrying the crRNA targeting A26) was incubated for 1 h at 70 °C in presence of 2.5 μM of Sso2081 dimer and various concentrations of target A26 ssRNA in a reaction containing 20 mM MES pH 6.0, 100 mM NaCl, 1 mg/ml BSA, 2 mM MgCl_2_, 0.5 mM ATP and a 5’-labeled A1 ssRNA that is not recognised by the Csm complex (5’-AGGGUAUUAUUUGUUUGUUUCUUCUAAACUAUAAGCUAGUUCUGGAGA). After 1 h incubation, 500 nM of Csx1 dimer was added to the reaction for a further 20 min. Reactions were quenched by deproteination with phenol-chloroform extraction and run on a 20% acrylamide 7 M urea denaturing gel prior to phosphorimaging to visualise A1 RNA cleavage.

### Generation of standards using the MazF nuclease

The *E. coli* toxin-antitoxin MazEF was purified as previously described ^10^. Active MazF was liberated by either trypsin (Promega) digestion (1600:1) at 37 °C for 15 min or by incubation with Factor X (0.1 unit per 1 mg of protein; Sigma-Aldrich) activated in FXa buffer containing 10 mM Tris-HCl pH 8.0 and 1 mM DTT. For generating linear oligoadenylates A_2_>P (5’ hydroxyl-Ap-Ap with cyclic 2’,3’ phosphate) and A_4_>P, 30 μM A_2_ (AAACAUCAG) or A_4_ (AAAAACAUCAG) MazF RNA was incubated with MazF in FXa buffer for 1 h at 37 °C. RNA was deproteinised by phenol-chloroform extraction followed by chloroform extraction. For use as standards, A_2_>P and A_4_>P linear oligoadenylates were 5’-end labelled using^32^P-γ-ATP and T4 Polynucleotide Kinase (PNK; Thermo Fisher Scientific) via its forward reaction.

### Thin-layer chromatography

Experiments were performed using silica gel on a 20 × 20 cm TLC plate (Supelco Sigma-Aldrich) containing a fluorescent indicator, allowing visualisation of non-radioactive products by UV shadowing. Prior to loading, all samples were deproteinised by phenol-chloroform followed by chloroform extraction. Non-radioactive or radioactive samples (from 0.1 μl to 2 μl) were loaded 1 cm above the bottom of the TLC plate. The TLC plate was then placed in a sealed glass chamber pre-warmed at 30 °C and containing 0.5 cm of a running buffer composed of H2O (30 %), ethanol (70 %) and ammonium bicarbonate (0.2 M), pH 9.3. The buffer was allowed to rise along the plate through capillary action until the migration front reached 15 cm. The plate was dried and radioactive sample migration visualised by phosphorimaging while non-radioactive sample migration was pictured under UV light (254 nm).

### Liquid chromatography-high resolution mass spectrometry (LC-HRMS)

LC-HRMS analysis was performed on a Thermo Scientific Velos Pro instrument equipped with HESI source and Dionex UltiMate 3000 chromatography system. Endlabeling of substrates or products was omitted. Samples were deproteinised as described for TLC. Compounds were separated on a Kinetex 2.6 μm EVO C18 column (2.1 × 100 mm, Phenomenex) using a linear gradient of acetonitrile (B) against 20 mM ammonium bicarbonate (A): 0 – 5 min 2 % B, 5 – 33 min 2 – 15 % B, 33 – 35 min 15 – 98 % B, 35 – 40 min 98 % B, 40 – 41 min 98 – 2 % B, 41 – 45 min 2 % B. The flow rate was 350 μl min^-1^ and column temperature was 40°C. UV data were recorded at 254 nm. Mass data were acquired on the FT mass analyzer in negative ion mode with scan range *m/z* 150 – 1500 at a resolution of 30,000. Source voltage was set to 3.5 kV, capillary temperature was 350 °C, and source heater temperature was 250 °C. Data were analysed using Xcalibur (Thermo Scientific). Extracted ion chromatograms were smoothed using the Boxcar function at default settings.

